# Peripheral opioid tolerance involves skin keratinocytes and platelet-derived growth factor type B signaling

**DOI:** 10.1101/2024.05.14.594040

**Authors:** Luca Posa, Sophia A. Miracle, Kathryn M. Albers, Matthew Fanelli, Angelique Buton, Timmy Le, Anita Khasnavis, Gilles Martin, Ashley K. McDonald, Mackenzie Gamble, Salome Fabri-Ruiz, Ryan W. Logan, Zachary Freyberg, Stephanie Puig

## Abstract

Opioid analgesic tolerance drives dose escalation which hampers the therapeutic utility of opioids by increasing centrally mediated deleterious side-effects, including respiratory depression or addiction. Peripheral opioid delivery provides a safer, effective alternative to systemic delivery by avoiding centrally mediated opioid side-effects. However, tolerance still occurs peripherally via mechanisms that remain unknown. Centrally, activation of the mu-opioid receptor (MOPr) by opioids induces release of platelet-derived growth factor-B (PDGF-B); and inhibition of PDGF receptor beta (PDGFRβ) prevents opioid tolerance. In the periphery, MOPr and PDGF-B are expressed in keratinocytes, and PDGFRβ is expressed in peripheral sensory neurons (PSNs), which are involved in tolerance. Previous studies showed that optogenetic stimulation of keratinocytes modulates PSNs via release of keratinocyte-derived factors. Thus, we hypothesized that mechanisms of peripheral opioid tolerance involve keratinocytes and PDGFRβ signaling. Using behavioral pharmacology, optogenetics and imaging in mice, we found that selective inhibition of peripheral PDGFRβ prevents peripheral morphine tolerance caused by repeated intraplantar (i.pl.) morphine injections. In addition, we show that PDGF-B is both necessary and sufficient to cause peripheral morphine tolerance. Repeated peripheral morphine injections lead to an increase in PDGF-B mRNA in MOPr-expressing keratinocytes and induce changes in the biophysical properties of keratinocytes as measured by patch-clamp electrophysiology. In parallel, we discovered that repeated optogenetic activation of keratinocytes is sufficient to induce peripheral morphine tolerance in a PDGF-B/PDGFRβ-dependent manner. Together, we show a novel epithelial-neuronal communication mechanism that incorporates keratinocytes and PDGF-B/PDGFRβ to mediate peripheral opioid tolerance, opening the door to safer, more effective pain therapeutics.

**Significance Statement:** Peripheral opioids are a safer alternative to systemic opioids. However, peripheral tolerance leads to reduction of analgesia over time, hampering clinical use of peripheral opioids. Here, we highlight a novel epithelial-neuronal communication mechanism that mediates peripheral tolerance. We discovered that intraplantar (i.pl.) morphine injections in mice cause peripheral tolerance via release of platelet-derived growth factor type B (PDGF-B) and activation of platelet-derived growth factor beta (PDGFRβ). We find that morphine i.pl. increases PDGF-B in mu-opioid receptor-expressing keratinocytes, which could be released to activate PDGFRβ in cutaneous nociceptor endings to mediate peripheral tolerance. Moreover, we show that photostimulation of keratinocytes is sufficient to cause peripheral tolerance in a PDGF-B/PDGFRβ-manner. Thus, keratinocytes and PDGF-B are new promising targets for peripheral opioid tolerance.

## Introduction

Opioids are powerful analgesic compounds that have long served as a cornerstone treatment for severe pain (1). However, repeated opioid use leads to tolerance, which requires escalating doses to overcome the reduction of analgesia over time. Dose-escalation reduces opioid safety by increasing deleterious centrally-mediated side effects such as physical dependence, respiratory depression, and/or addiction, which can lead to overdoses and death. Novel strategies to increase opioid safety without affecting analgesia are needed. Clinical evidence shows that peripheral and topical application of low doses of opioids produces effective analgesia, while limiting central penetration(2–5) and adverse side effects(3, 6–9). Peripheral and topical opioid formulations are in phase III clinical trials(10, 11), yet specific mechanisms of action for peripheral opioids remain poorly understood(12–14). In addition, even with peripherally administered opioids, tolerance to analgesia develops(14–18), overshadowing their clinical utility (2, 5, 19, 20).

Centrally, opioid activation of μ-opioid receptors (MOPr) causes spinal release of platelet-derived growth factor B (PDGF-B), which activates platelet-derived growth factor receptor beta (PDGFRβ) to directly cause spinal opioid tolerance[(21). Accordingly, imatinib, a PDGFRβ inhibitor, completely blocks spinal opioid tolerance(21), making PDGFRβ a promising target to prevent CNS tolerance. We therefore investigated whether PDGFRβ could also be an effective target to prevent peripheral opioid tolerance. In the periphery, MOPr, PDGFRβ and the PDGF-B ligand are expressed in somas of primary sensory neurons (PSNs) of dorsal root ganglia (DRG)(22–26). MOPr and PDGFRβ are also located in PSN nerve endings in the skin (25). Skin keratinocytes similarly express MOPr(24, 27–30) and PDGF-B (31–35). Skin MOPr activation impacts wound healing, inflammation and itch(28, 29, 36) but its role in tolerance remains unclear(37, 38). The convergence of MOPr and PDGF-B/PDGFR-β localization in peripheral tissue prompted us to hypothesize that PDGF-B/PDGFR-β signaling could have a role in peripheral opioid tolerance (39). In addition, the direct involvement of keratinocytes in somatosensation(40–43) raises the possibility that they are a component of the circuitry mediating peripheral tolerance. Anatomical and functional evidence indicate that keratinocytes communicate with PSN endings to modulate somatosensory information through keratinocyte-derived factors(40, 42, 44–47).

In this study, we used behavioral and pharmacological approaches to examine the role of PDGF-B/PDGFR-β signaling in opioid tolerance developed in response to peripheral morphine administration in mice. The impact of peripheral morphine administration and keratinocyte stimulation on expression of MOPr and PDGF-B in keratinocytes was examined using RNAscope fluorescent *in situ* hybridization. Patch-clamp electrophysiological recording on primary cultures of keratinocytes was used to examine the impact of repeated peripheral morphine administration on keratinocytes biophysical properties. Finally, targeted optogenetics was also used to determine the effect of keratinocyte activation on development of peripheral morphine analgesia and tolerance. We discovered that morphine peripheral tolerance is blocked by selective inhibition of PDGFRβ or PDGF-B in the periphery. Data also revealed that PDGFRβ and keratinocyte signaling are sufficient to induce peripheral morphine tolerance. In addition, peripheral morphine tolerance is accompanied by alteration of biophysical properties of keratinocytes as measured by patch-clamp electrophysiology. Thus, keratinocytes are an integral component of the circuit mediating peripheral tolerance, where activation of keratinocyte MOPr may evoke release of PDGF-B to activate PDGFRβ signaling and mediate peripheral morphine tolerance.

## Results

### Inhibition of PDGFRβ prevents peripheral morphine tolerance

We first established a model of peripheral morphine tolerance (**Fig. 1A**) by identifying a dose of intraplantar (i.pl.) morphine that would induce analgesia in the ipsilateral, but not the contralateral paw (N=6 males and females, 2-way ANOVA, Time x Paw, F (2, 20) = 8.954, P=0.0017, **Fig.1B, Sup. Table 1**). To test the impact of the PDGFRβ inhibitor, imatinib, on peripheral morphine tolerance, female (**Fig.1C left**) and male (**Fig.1C right**) mice received either vehicle (5μl), morphine alone (5μg/5μl), imatinib alone (10μg/5μl), or the combination of morphine + imatinib i.pl. for 5 consecutive days. Paw withdrawal latencies (PWL) were measured 20 minutes later to test analgesia and development of peripheral tolerance. Groups receiving morphine only developed peripheral morphine tolerance as shown by PWLs returning to baseline levels after 3 days of injections in both sexes (2-way ANOVA **Sup. Table 1**., **Fig. 1C left**: females, N=6, time x treatment, F (15, 100) = 9.411, P<0.0001; **Fig.1D right:** males, N=6, time x treatment, F (15, 100) = 4.917, P<0.0001). However, co-administration of morphine + imatinib i.pl. completely prevented tolerance, demonstrating that imatinib inhibition is sufficient to block peripheral morphine tolerance. A 3-way ANOVA analysis comparing sex x treatment x time showed no interaction (F (15, 246) = 0.808, P = 0.669), revealing no differential sex effect of imatinib on peripheral tolerance (**Fig. 1C, Sup. Table 1**.). We then combined data from both females and males and confirmed that imatinib prevented morphine tolerance (2-way ANOVA **Sup. Table 1., Fig. 1D**: N=12, time x treatment, F (15, 220) = 13.22, P<0.0001).

**Figure 1.**
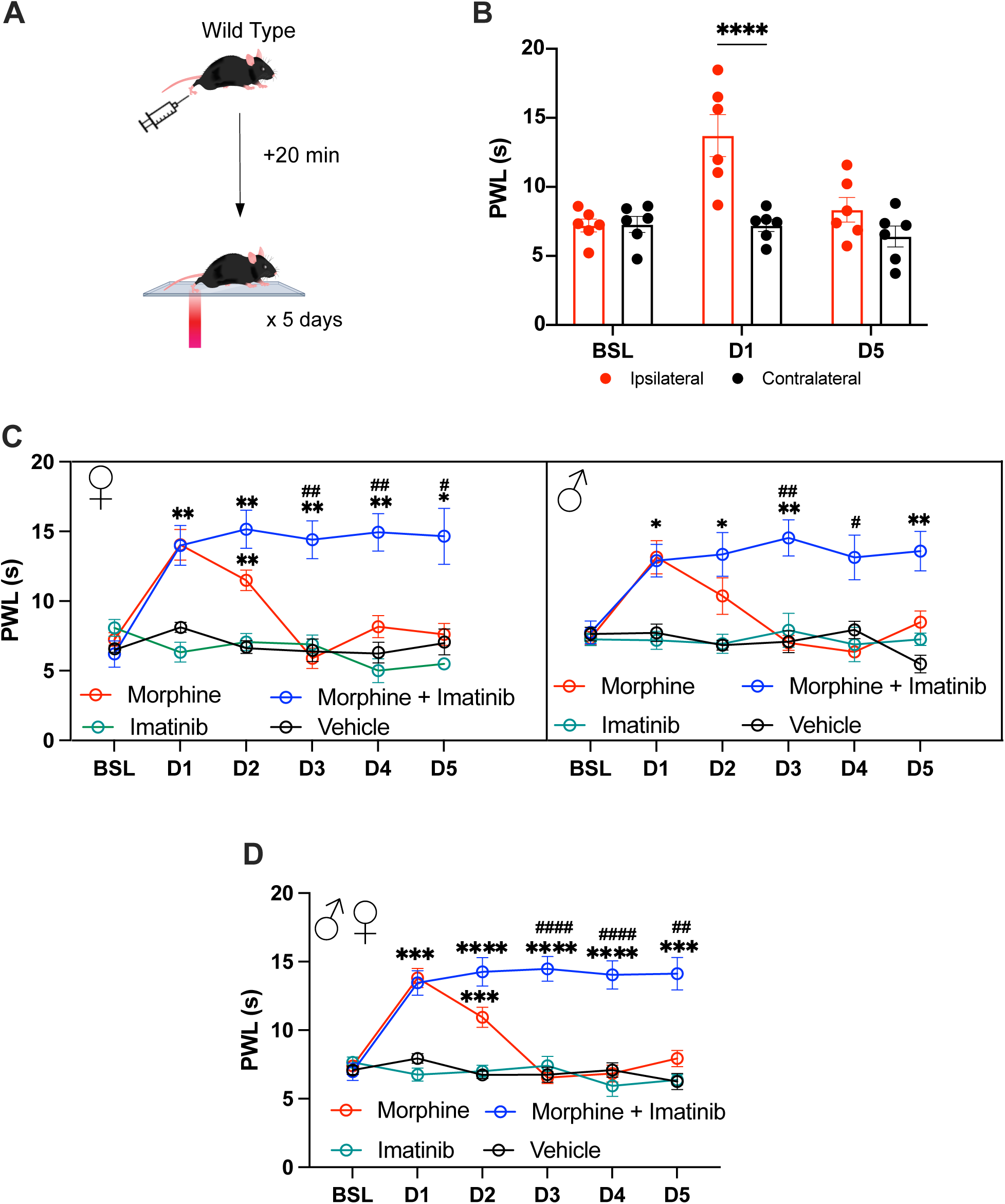
Imatinib blocks peripheral morphine tolerance. **A.** Schematic of the experimental design. **B**. Intraplantar (i.pl.) morphine administration induces ipsilateral but not contralateral acute antinociception as seen by increase in paw withdrawal latency (PWL) on day 1. Tolerance develops by day5. **C.** Morphine peripheral tolerance develops upon repeated morphine i.pl. injections. Co-injection of morphine + imatinib blocks peripheral tolerance in females (**C left**) and males (**C left**) mice. **D**. Females and males’ data merged. BSL = baseline, D = day. N = 6 per group/sex. * p<0.05, **p<0.01, ***p<0.001, ****p<0.0001 vs vehicle, # p<0.05 ## p<0.01, #### p<0.0001 vs morphine. Two-way or three-way Repeated Measures ANOVA followed by Šídák’s (**B**) or Tukey’s (**C-D**) multiple comparisons test. Data are expressed as mean ± s.e.m. Detailed statistics information can be found in **Sup. Table 1**.

### Administration of anti-PDGFRβ mAb blocks peripheral morphine tolerance

Though imatinib is a well-established PDGFRβ inhibitor, it may also target other receptor tyrosine kinases (48–54). Therefore, to thoroughly examine specificity of PDGFRβ inhibition on peripheral tolerance, we used a neutralizing monoclonal antibody (mAb), anti-PDGFRβ. When morphine was co-administered with PDGFRβ mAb for 5 days, peripheral morphine tolerance was completely attenuated in female and male mice (N=4-5 per sex per group, 2-way ANOVA, **Sup. Table 2**, **Fig. 2A left:** females, time x treatment, F (15, 70) = 9.797, P<0.0001; **Fig. 2A right**: males, time x treatment, F (15, 70) = 16.33, P<0.0001). No sex difference was found, as confirmed by 3-way ANOVA analysis (N=4-5/sex/treatment, F (15, 168) = 1.376, P = 0.164, **Sup. Table 2**). Importantly, no change in thermal pain threshold was observed over time in the groups treated with the mAb alone, showing that: 1) mAb PDGFRβ selective inhibition does not induce analgesia, and 2) the effect observed in combination with morphine specifically blocks peripheral morphine tolerance. Merging of female and male data confirmed that PDGFRβ mAb blocked morphine tolerance (2-way ANOVA **Sup. Table 1., Fig. 2B**: N=8-10, time x treatment, F (15, 160) = 22.82, P<0.0001). To further confirm that the i.pl. presence of an IgG2a isotype was not mediating these effects, we co-administered morphine i.pl. with IgG2a kappa isotype control antibody and found that it did not affect peripheral morphine acute analgesia and development of peripheral tolerance (**Fig. 2C**, N=4-9, 2-way ANOVA, time x treatment, F (10, 90) = 19.22, P<0.0001, **Sup. Table 2**). Collectively, these results support a specific role for PDGFR-β in mediating peripheral morphine tolerance.

**Figure 2.**
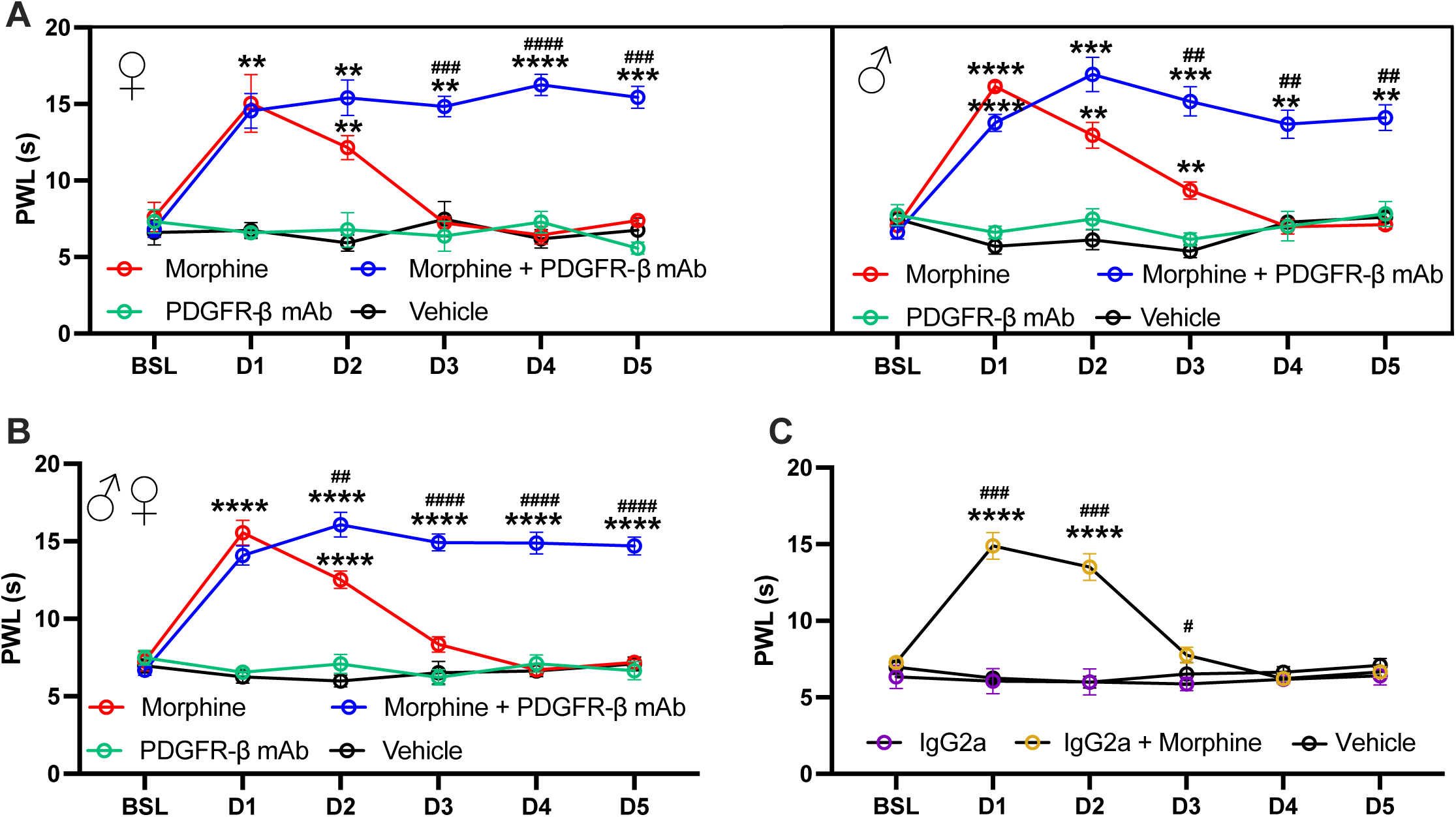
Selective PDGFRβ inhibition blocks peripheral morphine tolerance. Co-injection of morphine + anti-PDGFRβ monoclonal antibody (mAb) blocks peripheral tolerance in females (**A left**) and males (**A right**) mice. **B**. Females and males’ data merged. **C.** Control experiment. Neutral IgG2a mAb does not affect morphine acute analgesia on day 1 and does not prevent the development of tolerance overtime. BSL = baseline, D = day. N = 4-5 per group/sex. **p<0.01, ***p<0.001, ****p<0.0001 vs vehicle, # p<0.05 ## p<0.01, ### p<0.001 #### p<0.0001 vs morphine. Two-way or three-way Repeated Measures ANOVA followed by Tukey’s multiple comparisons test. Data are expressed as mean ± s.e.m. Detailed statistics information can be found in **Sup. Table 2**.

### PDGF-B is necessary and sufficient to cause peripheral tolerance

We next examined whether the endogenous PDGFR-β ligand, PDGF-B, is necessary for establishment of peripheral morphine tolerance. The PDGFR-β-Fc chimera protein (R&D Systems), which consists of a fusion of the N-terminal ligand binding domain of PDGFR-β with the Fc fragment of an IgG_1_, was used to confer solubility. This construct acts as a scavenger of extracellular PDGF-B, preventing its binding to PDGFRβ. Co-administration of morphine and PDGFRβ-Fc i.pl. completely abolished peripheral morphine tolerance in females (**Fig. 3A left,** N=6/group, 2-way ANOVA, time x treatment, F (15, 100) = 14.89, P<0.0001, **Sup. Table 3**), and males (**Fig. 3A right**, N=6/group, 2-way ANOVA, time x treatment, F (15, 100) = 13.44, P<0.0001, **Sup. Table 3**). These results show that extracellular PDGF-B at the periphery is necessary to cause peripheral morphine tolerance. A 3-way ANOVA analysis confirmed no sex difference between females and males (N=6/sex/treatment: F (15, 240) = 0.55, P**=**0.907**, Sup. Table 3).** Pooled data from both sexes further supports PDGFRβ-Fc as an effective inhibitor of morphine tolerance (2-way ANOVA; **Sup. Table 1**, **Fig. 3B**: N=12, time × treatment, F (15,220) = 28.01, P<0.0001).

**Figure 3.**
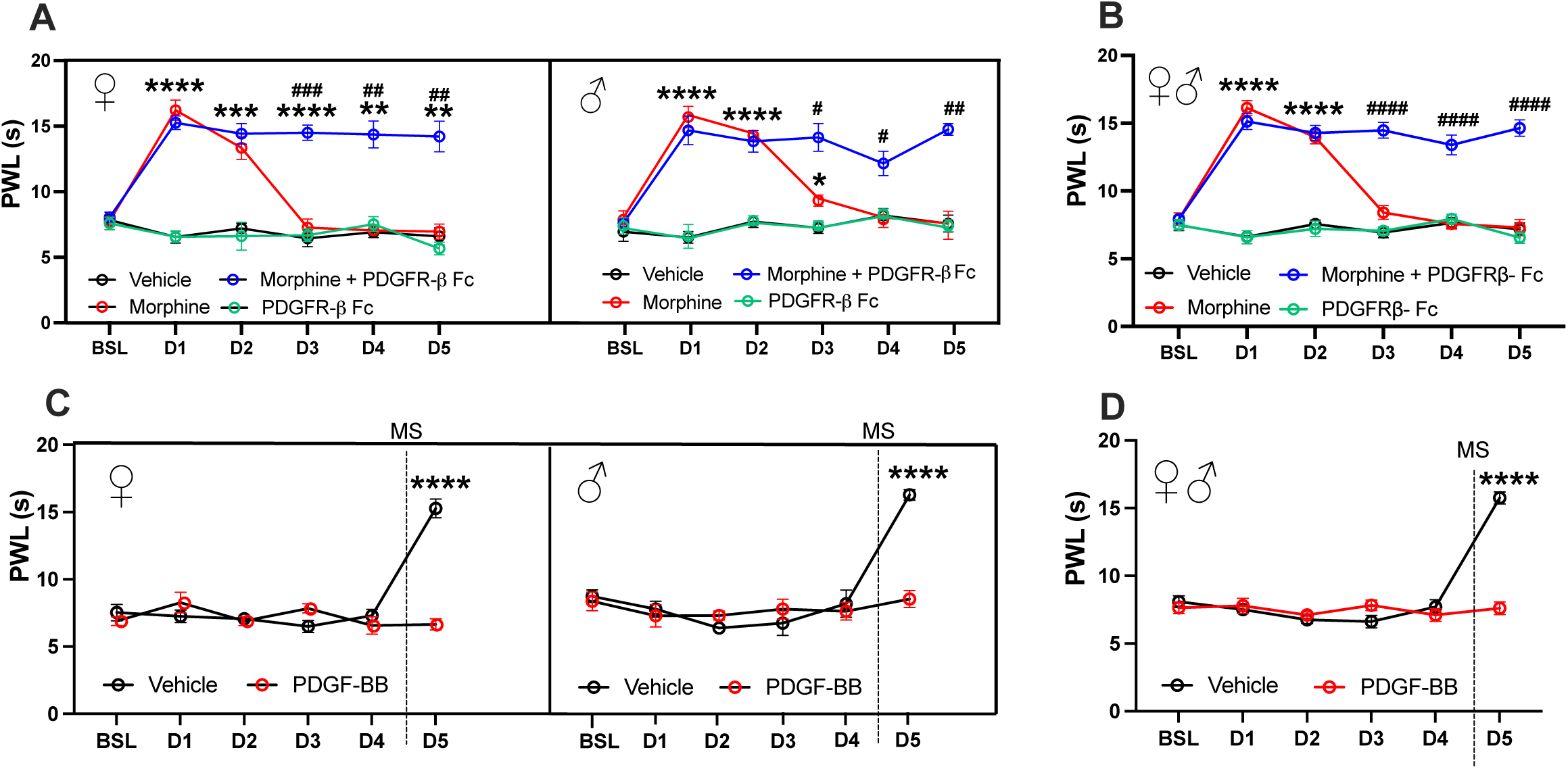
Selective PDGFRβ inhibition blocks peripheral morphine tolerance. A-C. Co-injection of morphine + the PDGF-B scavenger, PDGFRβ Fc, blocks peripheral tolerance in females (**A**) and males (**B**) mice. Morphine + PDGFR-β Fc females and males’ data merged (**C**). **D-F.** Repeated intraplanar administration of recombinant PDGF-B blocks analgesia to a morphine challenge on day 5, in female (**D**) and male (**E**) mice. Morphine + PDGFR-β Fc females and males’ data merged (**F**). MS = Morphine Sulfate, BSL = baseline, D = day. N = 5-6 per group/sex. **p<0.01, ****p<0.0001 vs vehicle, # p<0.05 ## p<0.01, ### p<0.001 #### p<0.0001 vs morphine. Two-way or three-way Repeated Measures ANOVA followed by Tukey’s (**A-B**) or Šídák’s (**C-D**) multiple comparisons test. Data are expressed as mean ± s.e.m. Detailed statistics information can be found in **Sup. Table 3**.

Next, we determined whether PDGF-B is sufficient to induce peripheral morphine tolerance. Vehicle or recombinant PDGF-B were administered i.pl. for 4 days. Consistent with prior work showing repeated spinal injection of PDGF-B did not alter baseline thermal thresholds, repeated i.pl. injection of PDGF-B did not change PWL thresholds in females (**Fig. 3C left**) and males (**Fig. 3C right**). On day 5, all mice were challenged with a dose of i.pl. morphine. Vehicle mice challenged with morphine showed acute antinociception. However, morphine antinociception was completely abolished in mice pre-treated with PDGF-B for 4 days in females (**Fig. 3C left**, N=6/group, 2-way ANOVA, time x treatment, F (5, 50)=26.72, P<0.0001, **Sup. Table 3**) and males (**Fig. 3C right**, N=5-6/group, 2-way ANOVA, time x treatment, F (5, 45)=11.43, **Sup. Table 3**). No interaction of sex x treatment was observed (N=5-6/sex/treatment, 3-way ANOVA, F (5, 114) = 0.609, P=0.693, **Sup. Table 3**). Combining data from both females and males confirmed that pre-treated with PDGF-B prevented acute morphine analgesia (2-way ANOVA **Sup. Table 1., Fig. 3D**: N=10-11, time x treatment, F (15,105) = 33.17, P<0.0001). These findings suggest that PDGF-B plays a pivotal role in peripheral morphine tolerance, as it is necessary and sufficient to induce morphine peripheral tolerance.

### Peripheral morphine administration increases PDGF-B expression in keratinocytes

We previously showed that PDGFR-β is expressed in cutaneous sensory neurons (25). PDGF-B ligand is also present in the periphery, with evidence of expression in epidermal keratinocytes (31–35). MOPr is similarly known to be expressed in DRG neurons (22, 23) and localized to terminals in the skin (24). Interestingly, MOPr mRNA (*Oprm1* gene) is also expressed in keratinocytes (24, 27–30). To examine coexpression of PDGFR-β, PDGF-B and MOPr, with a particular focus on keratinocytes, we used fluorescence in situ hybridization (RNAscope, ACDbio). In naïve mice, PDGF-B mRNA (*Pdgfb*) and *Oprm1* are co-expressed in keratinocytes, while PDGFR-β mRNA *Pdgfrb* was undetected (**Fig. 4A**). At baseline, 6.09+/-1.89% of total keratinocytes co-expressed *Pdgfb* and *Oprm1*(*Pdgfb+Oprm1+*), 5.68+/-1.73% only expressed *Pdgfb* (*Pdgfb+*), 6.079+/-2.11% only expressed *Oprm1*(*Oprm1+*), and 82+/-3.35% did not express either mRNAs (*Pdgfb-Oprm1-*, **Fig. 4B**). Interestingly, when compared to animals treated with morphine for 5 consecutive days and were peripherally tolerant to morphine, the number of *Pdgfb+* cells increased to 17.53+/-2.88%, and *Pdgfb+Oprm1+* cells increased to 27.4+/-3.09%, while *other* cells decreased to 47.14+/-3.59% (**Fig. 4B,** N=3-4/sex/group, 2-way ANOVA, Cell type x Treatment, F(3, 39)=32.87, P<0.0001, **Sup. Table 4**). Thus, the pool of cells expressing *Pdgfb* was increased by i.pl. morphine injections.

**Figure 4.**
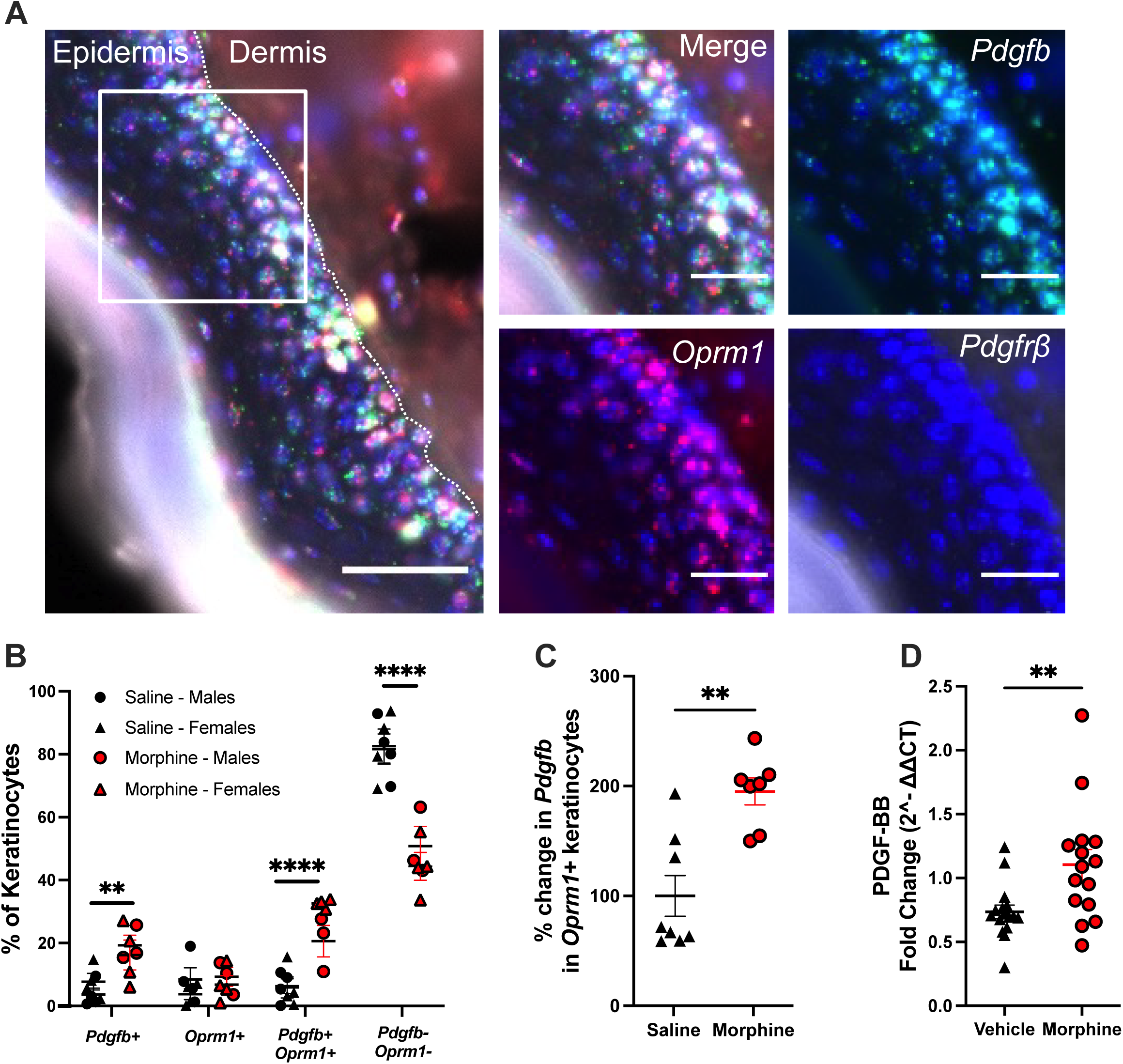
Peripheral morphine administration increases PDGF-B expression in keratinocytes. **A.** *In situ* hybridization shows co-expression of *Pdgfb* (green) and *Oprm1* (red) mRNA but not *Pdgfrb* (white) mRNA in epidermal keratinocytes. Scale bars: Left = 20μm, Zoomed-in figures = 10μm. **B.** Quantification of expression highlights that morphine i.pl. for 5 days changes proportions of keratinocytes expressing *Pdgfb, Oprm1,* or both. “Pdgfb+”: *Pdgfb* expressing cells; “Oprm1+”: *Oprm1* expressing cells; “Pdgfb+Oprm1+”: co-expressing cells; “Pdgfb-Oprm1-”: cells negative for both mRNAs. Two-way ANOVA followed by Sidack’s multiple comparison test. **C.** Amount of *Pdgfb* mRNA detected in *Oprm1* cells increases in mice injected with morphine i.pl. for 5 days. Unpaired Student t-test **D.** Total *Pdgfb* mRNA detected in glabrous skin with RT-PCR does not change in injected with morphine i.pl. for 5 days. Unpaired Student t-test. *Pdgfb* = PDGF-B mRNA, *Oprm1* = MOPR mRNA, *Pdgfrb* = PDGFRβ mRNA. **p<0.01, ****p<0.0001, morphine vs. vehicle. Data are expressed as mean ± s.e.m. Detailed statistics information can be found in **Sup. Table 4**.

Furthermore, the amount of *Pdgfb* in *Pdgfb+Oprm1+* keratinocytes was significantly increased in the morphine group compared to the saline group (**Fig. 4C**, Student unpaired 2-tailed t-test, P=0.0012, **Sup. Table 4**). This finding is also confirmed by an increase in total *Pdgfb* mRNA in skin samples from morphine i.pl. vs. vehicle i.pl. mice (**Fig. 4D**, Student unpaired 2-tailed t-test, P=0.0067, **Sup. Table 4**). These data indicate that repeated i.pl. morphine injections promote an increase in PDGF-B mRNA in keratinocytes that co-express both MOPr and PDGF-B.

### Peripheral morphine administration alters biophysical properties of keratinocytes

To examine whether repeated peripheral morphine administration could directly impact keratinocytes biophysical properties, we performed patch clamp electrophysiology on primary cultures of keratinocytes (primary KCs) from mice injected with morphine (MS-Tol) or vehicle (Veh) i.pl. for 5 days (**Fig. 5**). Importantly, we confirmed that *Pdgfb* and *Oprm1* mRNA were expressed in our primary KC culture using RNAscope (**Fig. 5A).** We showed that the input resistance of MS-Tol keratinocytes was significantly lower relative to their Veh counterparts (**; p=0.01, t=3.31, df=8; two-tailed unpaired Student’s t-test) (**Fig. 5B-E**). Additionally, the average resting membrane potential of keratinocytes in the Ms-Tol condition was significantly more negative (−27.75mV +/-10.69 mV) than that of Veh (7.29mV +/- 2.52 mV; p=0.003, t=3.97, df=9) (N=Vehicle: 6 cells; MS tol: 4 cells) (**Fig. 5B-D,F**). Moreover, voltage responses showed significant differences between the vehicle and morphine i.pl. primary KCs (**Fig. 5G**, p<0.001, **Sup. Table 5**). These data demonstrate that repeated peripheral morphine injections are impacting the overall biophysical properties of keratinocytes.

**Figure 5:**
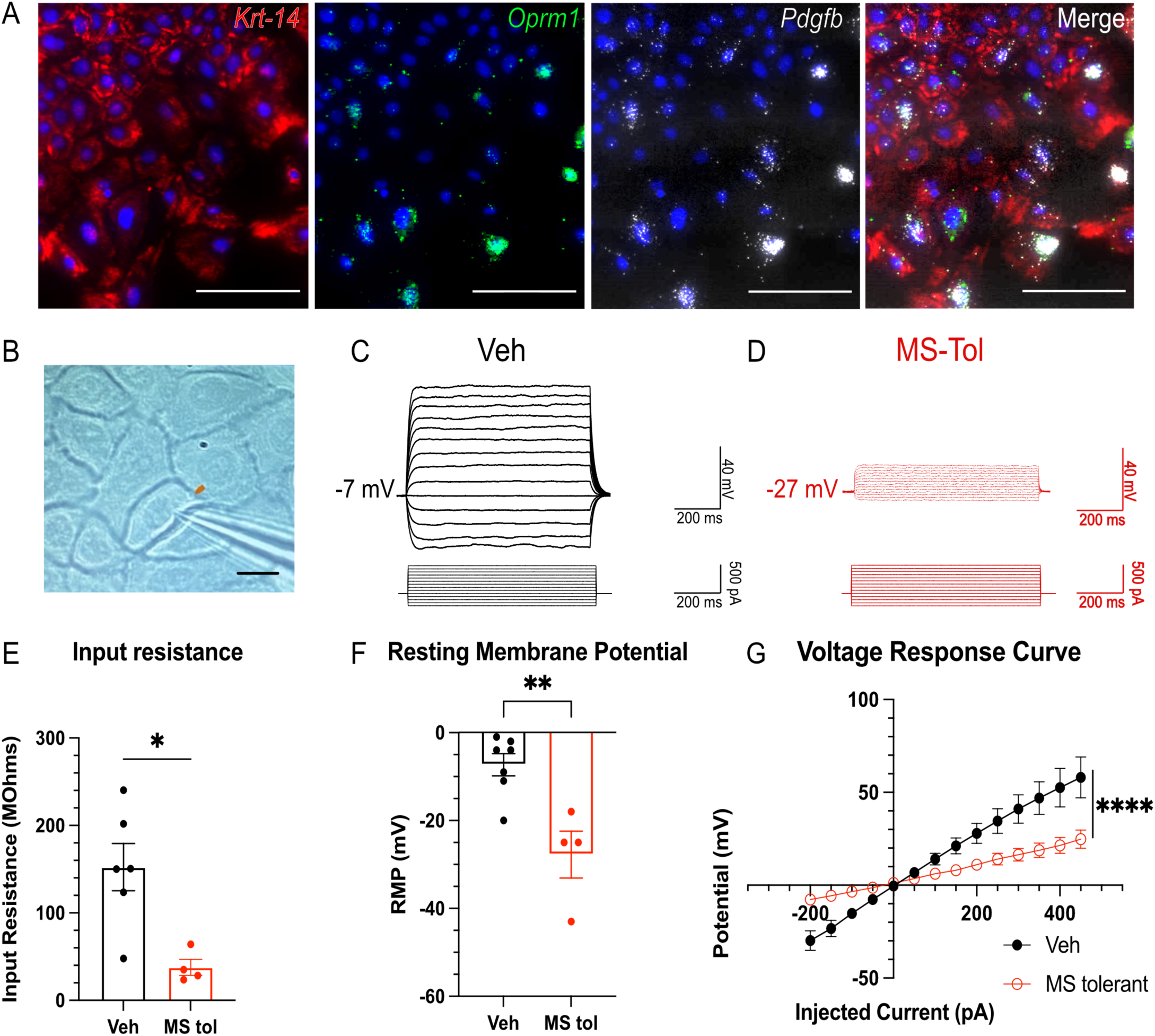
**Repeated morphine i.pl. injections decrease Input Resistance and Resting Membrane Potential in primary cultures of keratinocytes**. **A.** RNAscope highlighting expression of Krt14, Oprm1 mRNA and Pdgfb mRNA in primary culture of keratinocytes. **B.** Differential Interference Contrast (DIC) of primary cultured keratinocytes. Scale bar: 25 µm. Representative IV trace of **C.** Vehicle control and **D.** Morphine tolerant keratinocytes. Five days of morphine i.pl. decreases **E.** Input Resistance (two-tail unpaired Student’s t-test, p=0.01, t=3.31, df=8), and hyperpolarizes **F.** Resting Membrane Potential (two-tail unpaired Student’s t-test, p=0.003, t=3.97, df=9). F) in primary cultures of keratinocytes. **G.** Voltage response curve. (2-way ANOVA, p<0.0001) N=Vehicle: 6 cells; MS tol: 4 cells). *p<0.05 **p<0.01, ****p<0.0001. Detailed statistics information can be found in **Sup. Table 5**.

### Optogenetic stimulation of keratinocytes mediates peripheral morphine tolerance in a PDGFR-β- and PDGF-B-dependent manner

Evidence of PDGF-B and MOPr co-expression in keratinocytes and the necessity of PDGF-B signaling to mediate peripheral morphine tolerance suggested a mechanistic role for keratinocytes peripheral opioid tolerance. Earlier work showed that keratinocyte stimulation directly causes action potential firing in several PSN subtypes(40), suggesting functional communication between keratinocytes and PSNs, which are known to be involved in tolerance (37). We tested whether keratinocyte signaling is sufficient to drive opioid tolerance in the periphery using an optogenetic approach. In transgenic mice, blue-light sensitive channelrhodopsin-2 (ChR2) was genetically expressed in keratinocytes under the control of the Krt14 promoter (Krt14-ChR2 mice) (40). Krt14-ChR2 mice and wild type (WT) littermates were exposed to blue light for 5 consecutive days for 15min (BL-ON groups) (**Fig. 6A**). As a control, another cohort of Krt14-ChR2 and WT mice underwent the same procedure, but blue light was kept off (BL-OFF groups). On day 5, mice were injected with i.pl. morphine alone or in combination with imatinib (**Fig. 6B and C)**, or with PDGFRβ-Fc (**Fig. 6D and E**). Strikingly, the morphine analgesic effect was completely abolished in Krt14-ChR2 mice exposed to BL ON (**Fig. 6B morphine:** 3-way ANOVA, day x genotype x treatment, F (1, 24) = 6.014, P=0.0218, Sídák’s post hoc test BSL vs. Day 5: WT – morphine: P < 0.0001, K14Chr2 – morphine: P = 0.8438, **and 6D morphine:** 3-way ANOVA, day x genotype x treatment, F (1, 60) = 0.3537, P=0.5543, Sídák’s post hoc test BSL vs. Day 5: WT – morphine: P < 0.0001, K14Chr2 – morphine: P = 0.6315, **Sup. Table 5**). Importantly, challenge of morphine + imatinib to inhibit PDGFRβ (**Fig. 6B, morphine + imatinib**) or morphine + PDGFRβ-Fc (**Fig 6D, morphine + PDGFRβ-Fc**) produced a significant antinociceptive effect both in WT littermates and Krt14-ChR2 mice exposed to blue light (BL ON) for 5 days (**Fig 6B, morphine + imatinib**: Sídák’s post hoc test BSL vs. Day 5: WT – morphine + Imatinib: P < 0.0001, K14Chr2 – morphine + imatinib: P = 0.0003, **Fig 6D, morphine + PDGFRβ-Fc**: Sídák’s post hoc test BSL vs. Day 5: WT – morphine + PDGFRβ-Fc: P < 0.0001, K14Chr2 – morphine + PDGFRβ-Fc: P < 0.0001, **Sup. Table 6**). These data show that: 1) keratinocyte optogenetic activation ablates morphine analgesia; and 2) PDGFRβ signaling plays a crucial role in regulating these peripheral keratinocyte effects since morphine analgesia could be restored either by inhibition of PDGFRβ signaling via imatinib, or by scavenging PDGF-B with PDGFRβ-Fc. Importantly, control groups that underwent the same procedure but were not exposed to blue light (WT or Krt14-ChR2 BL-OFF) displayed the full analgesic effects of morphine when injected alone or in combination with imatinib (**Fig. 6C**, 3-way ANOVA, day x genotype x treatment, F (1, 16) = 0.1330, P=0.7201, Sídák’s post hoc test BSL vs. Day 5: WT – morphine: P =0.0002, K14Chr2 – morphine: P < 0.0001, WT – morphine + Imatinib: P < 0.0001, K14Chr2 – morphine + imatinib: P <0.0001, **Sup. Table 6**), or PDGFRβ-Fc (**Fig. 6E**, 3-way ANOVA, day x genotype x treatment, F (1, 40) = 3.892, P=0.0555, Sídák’s post hoc test BSL vs. Day 5: WT – morphine: P < 0.0001, K14Chr2 – morphine: P < 0.0001, WT – morphine + Imatinib: P < 0.0001, K14Chr2 – morphine + imatinib: P = 0.0011, **Sup. Table 6**). Altogether, these data demonstrate that repeated keratinocyte optogenetic activation is sufficient to induce peripheral morphine tolerance in a PDGFRβ / PDGF-B-dependent manner.

**Figure 6.**
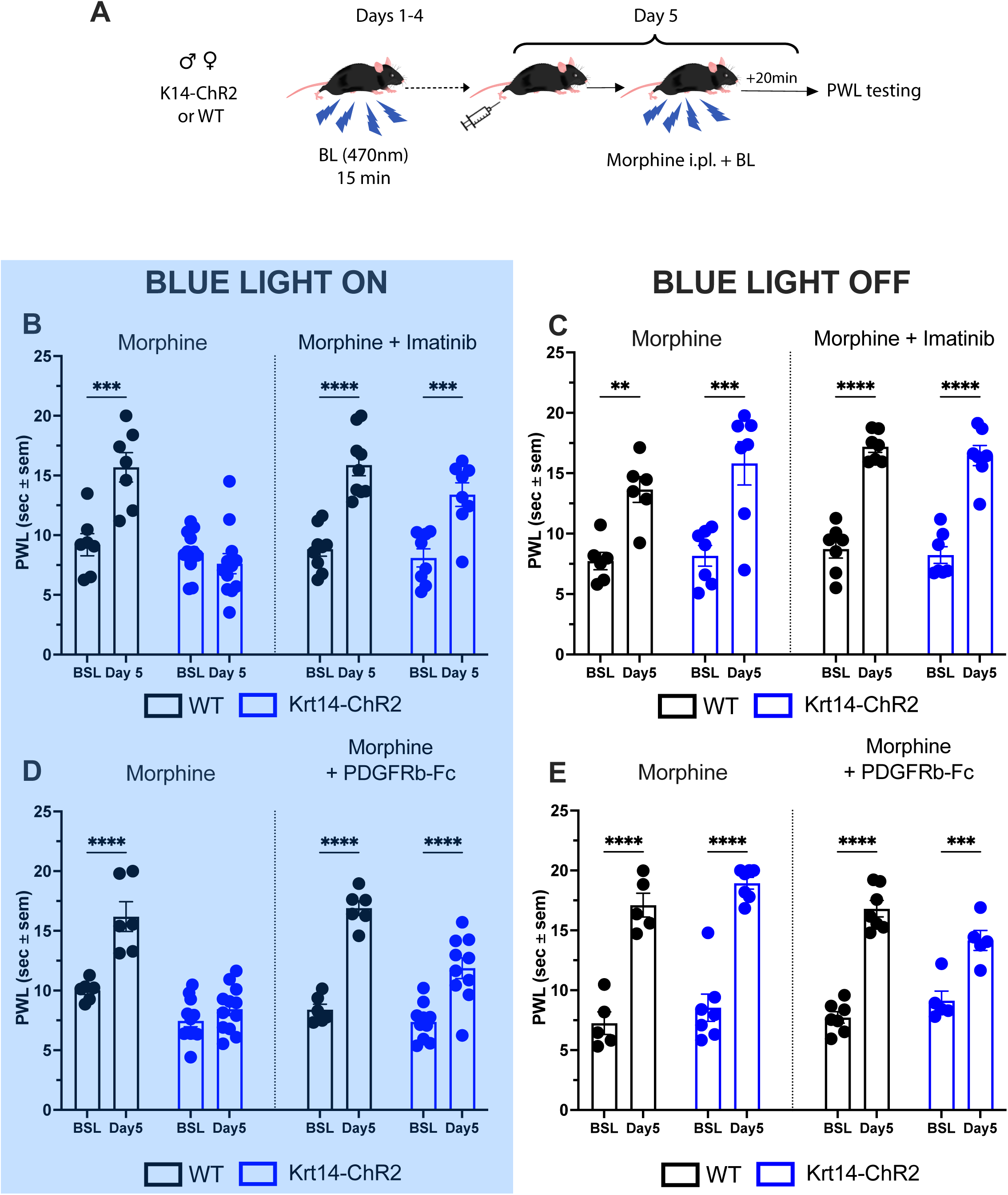
Optogenetic stimulation of keratinocytes mediates peripheral morphine tolerance in a PDGFR-β- and PDGF-B-dependent manner. **A**. Schematic of the experimental design. **B.** Blue-Light stimulation of keratinocytes for 5 days precludes the antinociceptive effect of a challenge of morphine on day 5, which is restored by a co-administration of morphine + imatinib. **D.** Blue-Light stimulation of keratinocytes for 5 days precludes the antinociceptive effect of a challenge of morphine on day 5, which is restored by a co-administration of morphine + the PDGF-B scavenger, PDGFRβ- Fc. **C, E.** Blue-light off controls. Three-way ANOVA followed by Sidak’s multiple comparisons test. **p<0.01, ***p<0.0001, ****p<0.0001. BSL vs. Day5. WT = Wild-Type. BSL = baseline. Data are expressed as mean ± s.e.m. Detailed statistics information can be found in **Sup. Table 6**.

### Optogenetic stimulation of keratinocytes does not increase PDGF-B expression

We examined the impact of optogenetic stimulation of keratinocytes in Krt14-ChR2 mice for 5 consecutive days, on *Pdgfb* expression and distribution in keratinocytes using *in situ* fluorescence hybridization (RNAscope, ACDbio, **Fig. 7A**). Intriguingly, and contrary to what was observed after 5 consecutive days of morphine i.pl. injections (**Fig. 7**), we did not observe a change in the population of cells expressing *Pdgfb, Oprm1* or both (**Fig. 7B,** N=3-4/sex/group, 2-way ANOVA, Cell type x Treatment, F(3, 42)=0.1550, P=0.9259, **Sup. Table 7**). The amount of *Pdgfb* in *Pdgfb+Oprm1+* keratinocytes or in bulk extraction of skin mRNA was not changed in the Blue-Light ON group compared to the Blue-Light OFF group (RNAscope: **Fig. 7C**, Student unpaired 2-tailed t-test, P=0.8109; RT-PCR: **Fig. 7D**, Student unpaired 2-tailed t-test, P=0.9866; **Sup. Table 7**). These data indicate that 15 minutes of daily optogenetic stimulation of keratinocytes is not sufficient to promote an increase in PDGF-B mRNA in keratinocytes.

**Figure 7.**
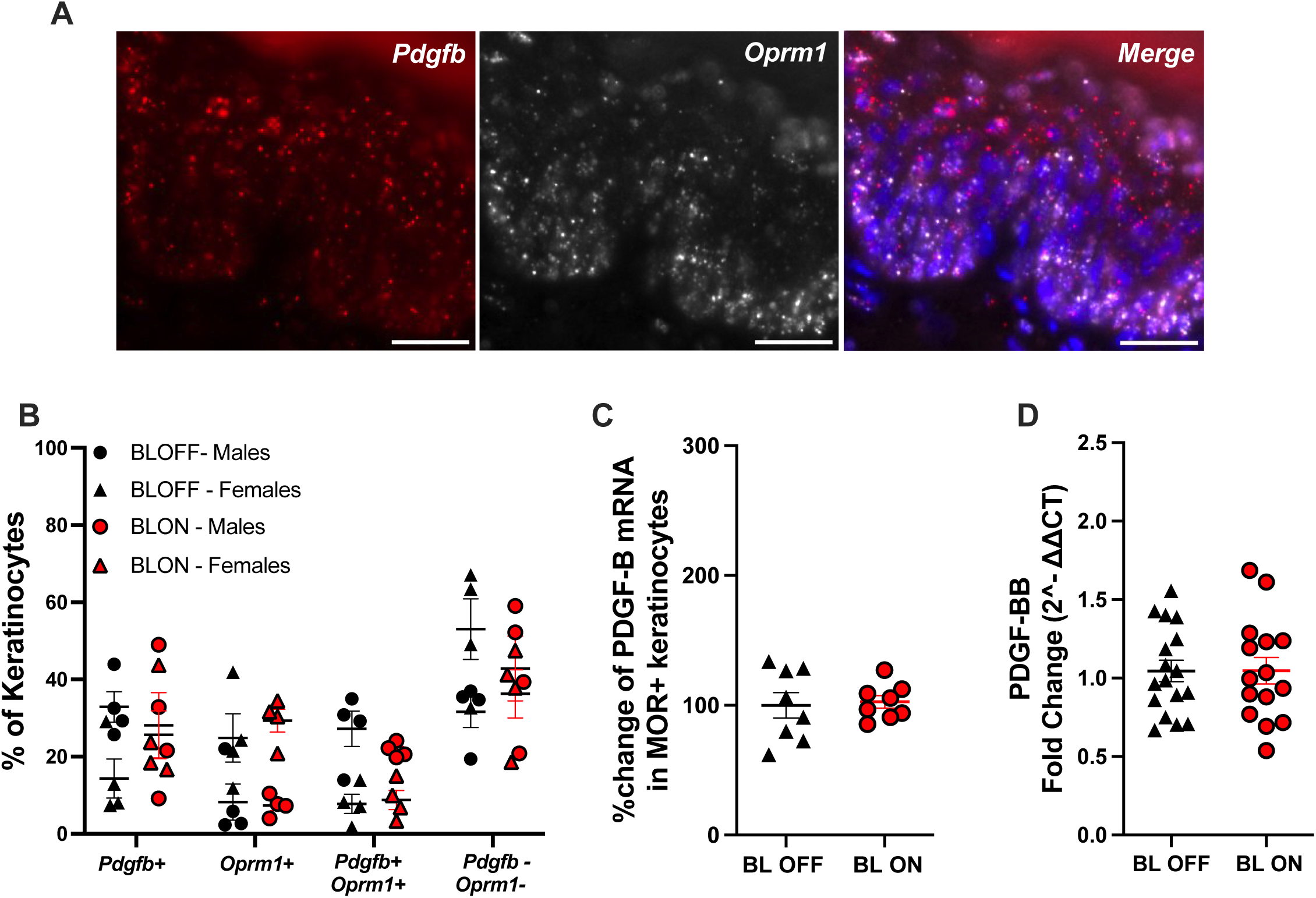
Keratinocyte photostimulation does not alter PDGF-B expression in the skin. **A.** *In situ* hybridization shows co-expression of *Pdgfb* (green) and *Oprm1* (red) mRNA in epidermal keratinocytes. Scale bars = 20μm. **B.** Quantification of expression highlights that 5 days of keratinocyte photostimulation in Krt14-ChR2 mice with blue light does not change proportions of keratinocytes expressing *Pdgfb, Oprm1,* or both. “Pdgfb+”: *Pdgfb* expressing cells; “Oprm1+”: *Oprm1* expressing cells; “Pdgfb+Oprm1+”: co-expressing cells; “Pdgfb-Oprm1-”: cells negative for both mRNAs. Two-way ANOVA followed by Sidack’s multiple comparison test. **C.** Amount of *Pdgfb* mRNA detected in *Oprm1* cells does not change in Krt14-ChR2 mice exposed to blue light, 15 min per day for 5 days. Unpaired Student t-test **D.** Total *Pdgfb* mRNA detected in glabrous skin with RT-PCR does not change in Krt14-ChR2 mice exposed to blue light, 15 min per day for 5 days. Unpaired Student t-test. BL ON = Blue Light On, BL OFF = Blue Light Off, *Pdgfb* = PDGF-B mRNA, *Oprm1* = MOPR mRNA. Data are expressed as mean ± s.e.m. Detailed statistics information can be found in **Sup. Table 7**.

## Discussion

In the current study, we investigated the involvement of PDGF-B/PDGFRβ signaling in peripheral morphine analgesic tolerance. We show that selective PDGFRβ inhibition completely blocks peripheral tolerance to morphine. Moreover, PDGF-B ligand is both necessary and sufficient to induce peripheral morphine tolerance. Furthermore, we demonstrate that skin keratinocytes express PDGF-B and MOPr, but not PDGFRβ, and repeated peripheral morphine administration increases the number of keratinocytes expressing PDGF-B, as well as overall PDGF-B expression in MOPr-expressing keratinocytes. Additionally, we show that repeated peripheral morphine administration causes profound changes in the biophysical properties of keratinocytes. Finally, using an optogenetic approach, we discovered that selective optogenetic stimulation of keratinocytes is sufficient to cause peripheral morphine tolerance in a PDGF-B/PDGFRβ-dependent manner.

This work is the first to investigate mechanisms of peripheral tolerance using a model employing peripheral morphine injections that induce analgesia locally. Importantly, our model enabled us to specifically investigate the mechanisms of peripheral tolerance without confounding factors of centrally-mediated MOPr signaling. This is important because prior studies suggested that peripheral tolerance mechanisms could be distinct from those involved in central tolerance(13, 55, 56). This assumption was supported by studies using topically applied opioids in animals that were tolerant to systemically administered opioids(56, 57). However, this idea was not confirmed by others(18, 58), leaving the issue of peripheral tolerance unresolved. Here, using our model of peripheral tolerance and a series of carefully designed pharmacology experiments, we provide evidence that peripheral tolerance shares common mechanisms with central tolerance through PDGF-B/PDGFRβ signaling. More specifically, we demonstrate that peripheral co-administration of a dose of morphine that only causes ipsilateral analgesia, with imatinib, a PDGFRβ inhibitor, completely blocks peripheral tolerance. Importantly, we confirm that peripheral morphine tolerance is also completely blocked with a selective PDGFRβ mAb, highlighting selectivity for PDGFRβ signaling in peripheral tolerance. Finally, we also show that a selective scavenger for extracellular PDGF-B (PDGFRβ-Fc) is also sufficient to preclude peripheral morphine tolerance.

Another important finding of our study is that selective *in vivo* photostimulation of keratinocytes using optogenetics in Krt14-ChR2 mice is sufficient to induce peripheral morphine tolerance, even in the absence of repeated opioid injections. This keratinocyte-mediated peripheral tolerance was precluded by acute co-administration of imatinib or PDGFRβ-Fc, indicating a previously unrecognized major role of keratinocytes/PDGFRβ signaling in peripheral tolerance. This startling finding highlights that repeated keratinocyte stimulation is sufficient to mimic the effects of repeated peripheral administration of morphine, and that both keratinocyte and morphine-mediated peripheral tolerance develop in a PDGF-B/PDGFRβ-dependent manner. This also provides the first evidence that keratinocytes are a component of the peripheral circuitry that conveys tolerance.

Involvement of keratinocytes in the peripheral circuits of tolerance is further supported by the fact that repeated morphine i.pl. profoundly alters keratinocytes biophysical properties as measured by patch-clamp electrophysiology. In control conditions, RPM and input resistance in these non-neuronal cells is consistent with prior work (59). We also found that repeated morphine exposure shifted keratinocytes’ resting membrane potential toward more negative values and decreased their membrane resistance. While the mechanism underpinning these changes remain unclear, the combined hyperpolarization and decrease in membrane resistance suggests that repeated morphine exposure leads to an increase of a potassium conductance. However, the G protein-coupled inwardly rectifying potassium channels that are activated by µ-opioid receptors in neurons are unlikely to account for the shift in resting potential in keratinocytes. Notwithstanding the fact that their activation depends on the presence of morphine, the KCNJ2 transcript coding for this channel is not expressed in keratinocytes (34). Additional studies are now needed to examine the precise mechanisms underlying peripheral morphine mediated changes in keratinocytes biophysical properties.

PDGFRβ is the receptor through which PDGF-B signals *in vivo*(60). Our study shows that PDGF-B/PDGFRβ signaling is both necessary and sufficient for peripheral morphine tolerance. In our prior work, we found that PDGFRβ is expressed in the soma and skin nerve endings of peripheral DRG neurons, specifically in small-diameter nociceptors, but not in larger myelinated neurons(25). In the current study, PDGFRβ mRNA expression was not found in keratinocytes, suggesting its involvement in peripheral tolerance is likely mediated by signaling in peripheral nociceptors. We also show that PDGF-B is expressed in mouse keratinocytes, consistent with single-cell transcriptomics analysis(34) and other studies of human and rodent epidermis assessing both PDGF-B mRNA and protein(33, 35, 61). Together, these data indicate that PDGF-B and PDGFRβ are present in peripheral nociceptive-conveying structures. In addition, 5 days of peripheral morphine administration increases PDGF-B expression in skin keratinocytes that co-express both MOPr and PDGF-B. This increased expression of PDGF-B in response to peripheral morphine administration, along with the blockade of peripheral tolerance by a PDGF-B inhibitor, collectively suggests that the peripheral release of PDGF-B upon opioid administration is necessary for peripheral tolerance and that keratinocytes may be the source of this growth factor. In fact, the concept that keratinocytes may be releasing PDGF-B upon opioid activation to initiate peripheral tolerance aligns with prior studies showing that keratinocytes can release factors to modulate peripheral nociceptors signaling activity(40, 42, 45, 47). Selective keratinocyte photostimulation using the same genetic construct as used in our study (Krt14-ChR2 mice) directly evoked action potentials in nociceptor-type PSNs when recorded at the level of DRGs(40). Importantly, the release of keratinocyte-derived factors(40, 47) is also thought to modulate peripheral sensory and nociceptive processing(42, 44–46, 62). However, we did not observe an increase of PDGF-B expression in keratinocytes after daily photo stimulation of keratinocytes despite precluding keratinocyte-mediated peripheral tolerance with the PDGF-B scavenger. We speculate that 15 min daily photostimulation of keratinocytes may not be sufficient to trigger changes in gene expression that are observed with repeated morphine i.pl. injections. However, our behavioral data indicates that this keratinocyte photostimulation was sufficient to induce peripheral morphine tolerance in a PDGF-B-dependent manner. We therefore speculate that keratinocyte photostimulation is sufficient to release PDGF-B to mediate the peripheral tolerance phenotype. Keratinocytes are known to release factors from vesicle content (47, 63, 64). Further experiments to directly examine mechanisms of PDGF-B release form keratinocytes and differences in mechanisms of release between morphine i.pl. and keratinocyte-mediated tolerance are underway.

Based on our data, we propose a model where opioid-activated keratinocytes engage PDGF-B/PDGFRβ signaling to cause peripheral tolerance by 1) releasing PDGF-B, and 2) activating PDGFRβ on peripheral nociceptors to mediate peripheral tolerance. This model also implies that MOPr signaling in keratinocytes is central to the initiation of peripheral opioid tolerance.

In summary, we demonstrate that PDGF-B/PDGFRβ signaling is an essential signaling mechanism involved in peripheral morphine tolerance. We found that keratinocyte stimulation is sufficient to induce peripheral opioid tolerance, highlighting that keratinocytes are an integrative component of the peripheral somatosensory circuit that mediates opioid signaling and peripheral tolerance. Our work opens new avenues of research on the potential of targeting keratinocytes and PDGFRβ to preserve peripheral morphine efficacy over time and enable the use of peripheral opioids for the treatment of local peripheral pain. If successful, this approach could be leveraged to shift opioid administration from central/systemic to localized/topical, thus circumventing the dangers of opioid addiction and abuse.

## Materials and Methods

### Animals

Mice were originally purchased from the Jackson laboratory (C57BL/6J (stock # 000664), Krt14-Cre (stock# 004782), Ai32 (stock #012569)). Animals were grouped and housed in a 12/12 light cycle (7 am lights on, 7 pm lights off). Water and rodent chow were provided *ad libitum* throughout the experiment. Animals were habituated to the laboratory environment for 1 week prior to experimental testing or breeding.

For optogenetic experiments, we generated Krt14-ChR2 mice by crossing mice that express Cre-recombinase under the control of the Krt14 promoter (Krt14-Cre) with mice that express the blue-light-sensitive channelrhodopsin-2 (ChR2) under the control of the CAG promoter inserted into the Rosa26 locus (Ai32) producing Krt14-ChR2 mice that express ChR2 in Krt14+ keratinocytes(40). F2 generation mice were used to obtain Cre allele heterozygous and either ChR2 heterozygous or homozygous mice. Littermate animals not expressing the ChR2 allele were used as wild-type (WT) controls. All experiments were performed on mice 8-14 weeks old.

All procedures were approved and performed in compliance with the Institutional Animal Care and Use Committee at Boston University and the University of Pittsburgh and University of Massachusetts Chan Medical School.

### Drugs

Morphine sulfate was obtained from Sigma (cat. number: 6211-15-0), Imatinib from LC Laboratories (Cat. # I-5508), recombinant rat PDGF-BB peptide (Cat #520-BB), recombinant mouse PDGFRβ-Fc scavenger (Cat. # 1042-PR) from R&D Systems, PDGFRβ monoclonal antibody (PDGFRβ mAb) from Invitrogen (Cat #16-1402-82), IgG2a kappa Isotype Control antibody from Invitrogen (Cat #16-4321-82). Drugs were either dissolved in a solution of filtered (0.22 μm) 10% β-cyclodextrin sulfobutyl ether (Captisol, CyDex) in 0.9% saline (for Imatinib, PDGFRβ-Fc, and PDGFRβ mAb), or in filtered 4mM HCl in 0.9% saline (for PDGF-BB peptide).

### Intraplantar (i.pl.) drug administration

All drugs were injected subcutaneously in the same region of plantar skin of one hind paw of awake animals that were lightly restrained. The volume of injection was 5μl using a 30-gauge needle and 20μl Hamilton syringe. The paw side was randomized within and across experiments. Doses per injection were as follows: Morphine Sulfate: 5μg, Imatinib: 10μg, PDGF-B peptide: 0.25μg, PDGFRβ-Fc: 250ng, PDGFRβ: mAb 0.1μg, IgG2: 0.1μg. Doses used were chosen after performing dose-responses (not shown).

### Nociceptive testing

#### Thermal Pain Assay

Mice were habituated to a Hargreaves apparatus (IITC Life Sciences Inc., Woodland Hills, CA) on a tempered glass maintained at 30°C for 90 minutes per day for 2 days and prior to behavioral testing. On test days, thermal nociceptive thresholds were recorded following habituation by measuring paw withdrawal latencies (PWL) to a light source focused on the plantar surface of each hind paw. Recorded behaviors for latency included withdrawal, licking, biting, or shaking of the targeted hind paw. Measurements were repeated three times per hind paw, with at least 1-minute intervals and averaged for each hind paw. Intensity of the targeted light source was adjusted to generate baseline responses at ∼8 seconds on average. A cut-off of 20 seconds was set to avoid tissue damage.

#### Experimental workflow

*Development of peripheral tolerance with morphine:* Mice were randomly assigned to drug groups. Development of tolerance and testing of pharmacological agents were performed in a 2×2 design by injecting either vehicle, morphine alone, the pharmacological agent alone, or morphine combined with the pharmacological agent. Mice were habituated to the Hargreaves apparatus for 60 to 90 min before drug injections. Baselines were acquired on day 0. On days 1-5, animals were injected intraplantarly (i.pl.) with an assigned drug, placed on the Hargreaves apparatus and PWL measured 20 minutes later. The 20-minute time point of testing was determined through a time course experiment to ascertain the peak analgesic effects of morphine administered i.pl. (not shown). On day 5, animals were euthanized and tissue collected according to tissue collection procedures.

#### Development of peripheral tolerance with blue light

On day 0, Krt14-ChR2 mice and WT littermates were habituated to the Hargreaves apparatus for 60 to 90 min before light exposure and drug injections. Following habituation, baseline PWL was measured using our Hargreaves apparatus. On days 1-4, animals were exposed to a blue light using a LED covered tray (170 Lumens, 475 nm wavelength) placed underneath the tempered glass surface. Blue light exposure lasted 15 minutes per day. On day 5, animals were injected in one hind paw (ipsilateral, ipsi) with either morphine or morphine combined with the tested agent, before being exposed to 15 min blue light exposure and paw withdrawal latency testing 5 minutes later. Control groups underwent the same procedure with the blue light off.

### Tissue Collection

Mice were anesthetized with 4% isoflurane vapor until respiration ceased followed by intracardiac perfusion. The abdomen was incised to expose the heart, a butterfly needle inserted into the left ventricle and the right atrium was cut to create an open circuit. Filtered 1X phosphate-buffered saline (PBS) was pumped through the circulatory system to remove all blood from the body. Hind paw skin was removed and cut in half longitudinally. One half was placed in 1mL of Trizol for RNA isolation and the other half was placed in 2% paraformaldehyde (PFA) at 4°C for 48 hours for fluorescence *in situ* hybridization analysis.

### Reverse Transcription - Polymerase Chain Reaction

For Reverse Transcription - Polymerase Chain Reaction analysis skin samples were homogenized in Trizol (Invitrogen, Cat. #15596026), incubated in 0.2mL chloroform for 2 min and centrifuged (15 min, 12,000 *xg* et 4°C) for phase separation. Directzol RNA isolation kits (Zymo Research, Direct-zol, cat #R2050) were used to isolate genomic DNA-free RNA. cDNA conversion was performed using the high-capacity RNA to cDNA kit (ThermoFisher, cat #438746). Input amount of RNA used was 2ug. cDNA reactions were done using a Thermocycler programmed at 37°C for 1 hour, 95C for 5min and held at 4°C. For RT-PCR reactions, 200ng in 2ul of sample per well were mixed with 5ul of TaqMan Gene expression MasterMix (ThermoFisher, Cat. # 4369016) and 2ul of H2O. The mix also contained 0.5ul of the following probes: Pdgfb (ThermoFisher, Assay ID: Mm00440677_m1) and housekeeping gene β-actin-FAM (ThermoFisher, Assay ID: Mm02619580_g1). No-RT controls were run side-by-side with samples. PCR was run using a Thermocycler programmed as follows: 1) 50°C for 2 minutes, 2) 95°C for 10 min, 3) 95°C for 15 secs, 4) 60°C for 1 minute. Steps 3 and 4 were repeated 40 times. Threshold cycles (Ct values) were used to calculate *Pdgfb* mRNA fold change. Briefly, ΔCt values were calculated by subtracting *Pdgfb*-Ct - *β-actin* Ct for each sample. ΔΔCt values were calculated by subtracting ΔCt values for each sample, to the average of ΔCt values of the control group for each experiment (Vehicle or BL OFF). Fold change was calculated with the following formula: 2^(-ΔΔCt).

### Fluorescence *in situ* hybridization

Skin tissue was transferred from 2% PFA to 30% sucrose for 24hrs, and then to 20% sucrose until sunk and stored at 4°C. Tissue was snap frozen in OCT, 14μm sections were obtained using a cryostat at −18°C and mounted on glass slides (FisherBrand™ Superfrost™ Plus Microscope Slides: 1255015). Sectioned tissue was stored at −80°C for at least 24 hours before use.

Slides stored at −80°C were transferred to an oven and baked at 60°C for 1 hour before post-fixation in 4% paraformaldehyde (PFA) for 1 hour at 4°C. After 2 washes in distilled water, slides were dehydrated in 50%, 70%, and 100% ethanol each for 5 mins. A hydrophobic pen was used to draw a barrier around tissue sections prior to incubation in target retrieval solution (ACDBio) at 95°C for 5 mins. Slides were then processed using the RNAscope kit V2 (ACDBio, Cat. 323100, Biotechne). Slides were incubated in hydrogen peroxide 10 min at RT and protease III at 40°C for 30 mins. Probes targeting mRNAs sequences of the platelet-derived growth factor type b (Pdgfb, Mm-PDGF-B-C1 (ACD Bio Cat. 42451)), platelet-derived growth factor receptor beta (Pdgfrβ, Mm-PDGFr-B-C2 (ACD Bio Cat. 411381-C2)) and opioid receptor mu-1 (Oprm1, Mm-Oprm1-C3 (ACD Bio Cat. 315841-C3)), were added to the slides and incubated at 40°C for 2hrs. Probe signals were then amplified using manufacturer’s AMP solutions 1, 2, and 3 via incubation at 40°C for 30 mins, 30 mins, and 15 mins, respectively. Probes were assigned a TSA Plus fluorophore (TSA Vivid 520 - Pdgfb, TSA vivd 570 - Pdgfrβ, TSA vivid 650 - Oprm1) and incubated for 30 min at 40°C. Slides were incubated with HRP blockers at 40°C for 15 mins following each TSA Plus fluorophore incubation. All tissue was stained with DAPI (ACD Bio Cat. 323108), coverslipped and slides stored at 4°C until imaging.

### RNAscope imaging and analysis

Skin tissue labelled with fluorescent mRNA probes was imaged using a BZ-8000 widefield fluorescence microscope (Keyence) using a 40X objective lens. Z-stacks were acquired (3μm total, 1μm steps). Two ROIs were imaged per tissue sample per mouse and Maximum Intensity projections were used for analysis.

All images were uploaded to QuPath for mRNA spots analysis. Cell segmentation was performed based on DAPI nuclear staining. The basal layer of keratinocytes was segmented using the following settings: 10μm background radius, Sigma 0.75, Minimum area: 2μm^2^, Maximum Area 400μm^2^, Cell expansion: 2μm. The upper layers of keratinocytes were segmented using the following settings: 8μm background radius, Sigma 0.75, Minimum area: 2μm^2^, Maximum Area 400μm^2^, Cell expansion: 4μm. Thresholds for detection of spots (subcellular detections, mRNA copies) were calculated as 35% of the maximum intensity of each channel in the ROI. Minimum and maximum spot size set at 0.3 and 2μm, respectively. Background noise was calculated by averaging the number of estimated spots per cell across all samples in an experiment. Cells were then considered positive for an mRNA if they contained at least the average detected spots. Cell populations were categorized as: 1) negative for both Pdgfb and Oprm1 mRNA (*Pdgfb-/Oprm1-),* negative for Pdgfb and positive for Oprm1 (Pdgfb-/Oprm1+), positive for Pdgfb and negative for Oprm1 (Pdgfb+/Oprm1-), and positive for Pdgfb and positive for Oprm1 (Pdgfb+/Oprm1+). Then, we calculated the average number of Pdgfb mRNA copies in all Oprm1+ cells per treatment group.

### Primary Cultures of Keratinocytes

Glabrous skin pads were collected from mice on day 5 after vehicle or morphine i.pl. treatments. They were rinsed in betadine, Hank’s Balanced Salt Solution without calcium or magnesium (HBSS; from Fisher Scientific cat # 14170112), then 70% EtOH (<1 minute per rinse). Skin pads were placed in Dispase solution (8mg/ml Dispase II [neutral protease, grade II from Sigma Aldrich cat# 4942078001] in HBSS with 0.1% sterile Penicillin-Streptomycin [P/S; 5,000U/ml from Fisher Scientific cat#15070063] and 0.5μg/ml Amphotericin B solution fungizone [from Sigma Aldrich cat# A2942]) for 1 hour at 37°C. Skin pads were agitated every 10 minutes during this incubation. The epidermis was then peeled from the dermis of each pad, and the epidermis was submerged in Trypsin-EDTA (0.05%, from Fisher Scientific cat#25300054) at 37°C for 30 minutes, agitating every 5 minutes. Next, equal volume of HBSS was added to the Trypsin-EDTA solution and the skin pads were mechanically ground with a pipette tip for further dissociation. This solution was passed through a 70um cell strainer, then was centrifuged at 0.6RCFx1000 for 5 minutes. The top layer was removed, and the cells pelleted at the bottom were resuspended in complete Defined Keratinocyte Serum Free Basal Medium (KSFM [from Fisher Scientific cat #10744019], with 0.2% defined growth supplement [from Fisher Scientific cat #10744019], 0.1% P/S, 100μg/ml Epidermal growth factor [EGF; from Fisher Scientific cat#53003-018], and 0.5μg/ml Amphotericin B solution fungizone). Cells were plated on collagen (from Fisher Scientific cat#NC1558174)-coated coverslips and allowed to grow at 37°C and 5% CO_2_ with KSFM media changes every other day. Keratinocytes were recorded using whole-cell patch clamp 4-7 days after plating.

### Electrophysiology

Whole-cell patch clamp recordings of keratinocytes were performed in a recording chamber filled with artificial cerebral spinal fluid (aCSF) of the following composition (mM): 126 NaCl, 2.5 KCl, 1.3 NaH_2_PO4×H_2_O, 1 MgCl_2_, 2 CaCl_2_, 26 NaHCO_3_, 10 D-Glucose, at a rate of 2-3 ml/min at room temperature. Briefly, borosilicate glass electrodes (1.5 mm OD, 7– 14 MΩ resistance) were filled with an internal solution containing (mM): 120 K-methanesulfonate; 20 KCl; 10 HEPES; 2 ATP, 1 GTP, and 12 phosphocreatine. Following seal rupture, series resistance was 29.2 ± 1.1 MΩ, fully compensated in current clamp recording mode, and periodically monitored throughout recording sessions. Recordings with changes of series resistance larger than 20% were rejected. Current-Voltage traces were recorded with the following parameters: 14 sweeps with increasing injected current of +50 pA with an initial injected current of −200 pA. Voltage and current traces in whole-cell patch-clamp were acquired with an EPC10 amplifier (HEKA Elektronik; Germany). Sampling was performed at 10 kHz and digitally filtered voltage and current traces were acquired with PatchMaster 2.15 (HEKA Elektronik; Germany) at 2 kHz. All traces were subsequently analyzed off-line with FitMaster 2.15 (HEKA Electronik; Germany). Experimenter was blinded during patch experiment and analysis but was unblinded during data interpretation.

### Data Analyses and Statistics

Behavioral and cell counting measures were performed by investigators blinded to experimental groups. All results are presented as mean ± s.e.m. Data were tested for normality and sphericity and a Greenhouse Geisser correction was applied where appropriate. Assumption of Normality was tested either with the Shapiro–Wilk test or with a Q-Q plot. Student unpaired 2-tailed t-test was used to compare means of two independent groups. Repeated Measure two-way or three-way ANOVA were used with treatment and time or treatment, time and sex as variables, respectively. Tukey’s or Šídák’s were used for *post hoc* comparisons of groups. GraphPad Prism 9.5 and RStudio and R software (R 4.3.1.) were used for data treatment and statistical analysis. All statistics are summarized in Supplementary Tables 1-4. P<0.05 was considered as significant.

## ACKNOWLEDGMENTS

We thank Ms. Bahhiyah Jefferson, research technician in the Albers laboratory at the University of Pittsburgh for her technical assistance with experiments for this project. We also thank Ms. Kennedy O’Hara, first year graduate student in the Morningside Graduate School at UMass Chan for her technical assistance with experiments for this project.

All data in this manuscript is available upon request to the corresponding author.

For this work, SP was supported by the National Institute of Health (NIH) R21DA051636. KMA was supported by NIH R21DA051636, R01AR0777341 and R01DK124955. RWL was supported by NIH R01DA051390 and R01DA06124. ZF was supported by The Pittsburgh Foundation (John F. and Nancy A. Emmerling Fund of the Pittsburgh Foundation, FPG00043), the Commonwealth of Pennsylvania (PA-HEALTH to Z.F.), and NIH R21AA028800; R01ES034037 and R01DA061243.

## Author Contributions

SP designed the study and obtained funding, KMA helped obtain funding, KMA and RWL helped design study; LP, SP, SAM, AKM MF, AK, TL, AB performed experiments; SP and LP; MG, KMA, ZF, RWL, GM and SFR helped with analysis; LP and SP wrote the manuscript.

## Project Funding

NIH, NIDA 1R21DA051636. PI: Stephanie Puig

